# Organisation of axial regions of isolated mitotic chromosomes visualised by cryo correlative light and electron tomography

**DOI:** 10.1101/2025.02.06.636497

**Authors:** Gurudatt Patra, Mohamad Harastani, Kumiko Samejima, Lucy Remnant, Nathalie Troffer-Charlier, Corrine Crucifix, Alexandre Durand, Nils Marechal, Yves Lutz, Anna M. Steyer, Zhengyi Yang, Wim J. H. Hagen, William C. Earnshaw, Mikhail Eltsov

## Abstract

The formation of mitotic chromosomes is essential for the accurate segregation of genetic material during cell division. Increasing evidence suggests that chromosome formation involves the reorganization of DNA into loops anchored within chromosomal axial regions, whose structural organization remains insufficiently characterized. Taking advantage of DT40 cells, an avian cell model characterized by the presence of a range of chromosome sizes from 3.2-197 Mb, we have established a preparation of entire close-to-native native mitotic chromosomes for cryogenic correlative light and electron microscopy (cryo-CLEM). The size of the smallest chromosomes allows imaging of their axial regions without further thinning. Cryo-electron tomography of the chromosome axial regions reveals the presence of heterogeneous non-histone macromolecular densities (NHMDs), approximately 30–45 nm in size, interspersed within chromatin/DNA regions. We propose that NHMDs may contain condensins and contribute to chromosome architecture. In addition to NHMDs, we identified dense clusters of particles, similar in size, near the chromosome surface, likely associated with ribosomal components. To quantitatively differentiate NHMDs from these surface clusters, we developed an analytical approach based on particle interspacing and spatial distribution within the chromosome volume. By establishing a cryo-CLEM workflow for whole, near-native mitotic chromosomes, our study provides a foundation for investigating their ultrastructural architecture.

## Introduction

The first level of DNA compaction in chromatin involves a “bead-on-a-string” arrangement, where DNA wraps around histone octamers forming nucleosomes connected by linker DNA (Luger et al. 1997), (Oudet, Gross-Bellard, and Chambon 1975), (Kornberg and Lorch 1999). During mitosis, eukaryotic chromatin undergoes further compaction which is critical for accurate segregation of genetic material between the daughter cells. However, the exact mechanism of how chromatin organizes into the highly compact rod-like mitotic chromosomes is still in question (Dietzel and Belmont 2001), (Swedlow and Hirano 2003), (Maeshima and Eltsov 2008). *In vitro* experiments initially carried out in the 1980s treated isolated mitotic chromosomes with polyanions or high salt to remove histones and revealed DNA loops arranged around a central proteinaceous structure made up of a network of non-histone proteins (J. R. Paulson and Laemmli 1977; Earnshaw and Laemmli 1983). These residual structures were named chromosome scaffolds because they resembled the size and shape of the original chromosomes when imaged in an electron microscope. It was hypothesized that the scaffolds were responsible for maintaining the chromosome shape and integrity. The major components of the scaffolds were found to be topoisomerase II and condensin complexes (Maeshima and Laemmli 2003), (James R. Paulson et al. 2021). The initial concept of a continuous proteinaceous chromosome scaffold was challenged by biochemical and microscopy studies (Okada and Comings 1980) and later by micromechanical force measurements (Poirier and Marko 2002) that demonstrated chromosome disintegration upon nuclease treatment. Thus, the existence and organization of chromosome scaffolds remain elusive.

Most eukaryotes have two types of condensin complexes, condensin I and condensin II that play an important role in chromosome condensation, essential for structuring mitotic chromosomes (Hirano 2012), (Hirano 2016), (Gibcus et al. 2018), (Green et al. 2012). Condensin I and II are multisubunit complexes that share a common pair of structural maintenance of chromosomes (SMC) ATPase subunits (SMC2 and SMC4), that bind to distinct sets of three regulatory subunits. *In vitro* studies have demonstrated DNA loop extrusion activity mediated by condensins (Ganji et al. 2018), (Golfier et al. 2020), (Ryu et al. 2022). Early immunofluorescence and immunogold labelling of condensins revealed their localization along the central axial region of both the sister chromatids of isolated chromosomes and also in chromosomes observed directly within mitotic cells (Hudson et al. 2003) (Maeshima, Eltsov, and Laemmli 2005a), (Ono et al. 2004).

While these axial condensin-enriched regions corresponded in principle to the EM observations of chromosome scaffold structures, the structural relevance of the chromosome scaffold has been questioned. Earlier studies on the chromosome scaffold that involved histone depletion from isolated chromosomes by heparin/dextran sulfate (Earnshaw and Laemmli 1983) or by high concentration of NaCl (Adolph et al. 1977) are harsh procedures that result in artificial rearrangements or even precipitation of non-histone components (Okada and Comings 1980). It should be noted that the original scaffold observations were done after the complete drying and contrasting of the preparations with heavy metal salts which could result in additional structural artefacts (Dubochet et al. 1988). In contrast, more recent studies using superresolution light microscopy without histone depletion or drying showed a less defined distribution of both condensin complexes in human mitotic chromosomes (Walther et al. 2018). While condensin II had more axial distribution, condensin I was spread within ∼50% of the chromatid diameter (Walther et al. 2018), (Green et al. 2012), (Gibcus et al. 2018).

Cryo-electron tomography (cryo-ET) of frozen-hydrated samples enables close-to-native three-dimensional imaging at molecular resolution by preserving biological samples in a vitrified state. Recently, this technique was applied to investigate isolated HeLa chromosomes vitrified on EM grids (Beel et al. 2021) and intact mitotic HeLa cells imaged *in situ* within cryo-lamellas obtained with cryo-focused ion beam milling (Chen et al. 2024). These studies provided information on nucleosome–nucleosome interactions and the local geometry of chromatin fibres. However axial non-histone components of chromosomes were not reported, likely because of technical limitations related to the localization of the axial areas and the high-crowding of the mitotic chromatin landscape. To address this knowledge gap, we developed a cryo-CLEM pipeline to target the chromosome axial regions for ultrastructural analysis. We leveraged the advantages of the chicken DT40 system, whose genome contains macro, mini and dot chromosomes (Z. Huang et al. 2023). Cell synchronization and chromosome isolation have been established for this system in previous studies (Earnshaw et al. 1985), (Lewis and Laemmli 1982) (Ohta et al. 2010). To reduce chromatin crowding and make the axial regions of chromosomes more accessible for cryo-ET imaging without additional thinning, we applied a partial decondensation by incubation with a low ionic strength buffer. Cryo-FIB milling was avoided due to the technical challenges associated with aligning lamellae along the chromosome axis. Instead, the axial regions were targeted using fluorescent labelling of halo-tagged SMC2, and localized in complete chromosomes by cryo-fluorescence imaging, followed by cryo-ET and computational denoising. This approach enabled visualization of nucleosomes, DNA linkers, and non-histone macromolecular densities (NHMDs) within the axial region of the mitotic chromosome, while aggregated densities likely to be ribosome-related were observed at the chromosome surface. NHMDs were found to exhibit a high structural heterogeneity and were interspersed with chromatin. This study represents an initial observation of metazoan mitotic non-histone components in the context of mitotic chromosomes and establishes a methodological foundation for their detailed structural characterization.

## Results

### Preparation of chicken DT-40 chromosomes for cryo-EM

Our chromosome preparation pipeline was developed to achieve the following objectives: (i) reproducible attachment and uniform distribution of chromosomes on cryo-EM grids; (ii) complete vitrification within a thin buffer layer during plunge freezing; and (iii) preservation of chromosome integrity with the axial region of the chromosome compatible for cryo-ET imaging.

We investigated the partial decondensation of chromosomes in HEN buffer (10 mM Hepes pH 7.5, 0.2 mM EDTA, 25 mM NaCl, 0.1% NP 40). This buffer leads to chromosome decondensation by washing out polyamines contained in the storage buffer while maintaining a physiological pH. 0.5 mM ATP ɣS was added to stabilize condensin binding to chromatin to minimise its dissociation during decondensation (Ganji et al. 2018), (Terakawa et al. 2017). A visible decondensation was observed (Figure 1A), as observed by fluorescence labelling of DNA by Hoechst 33342 (from Sigma-Aldrich, Catalog: B2261). We show below that one of the major axial proteins of the mitotic chromosome maintains its axial localization in this partially decondensed condition. The length of the major micro chromosome population visualized in our coverslips and grids was 1-3 μm in polyamine-containing storage buffer and 2-10 μm after decondensation in HEN buffers.

**Figure 1:**
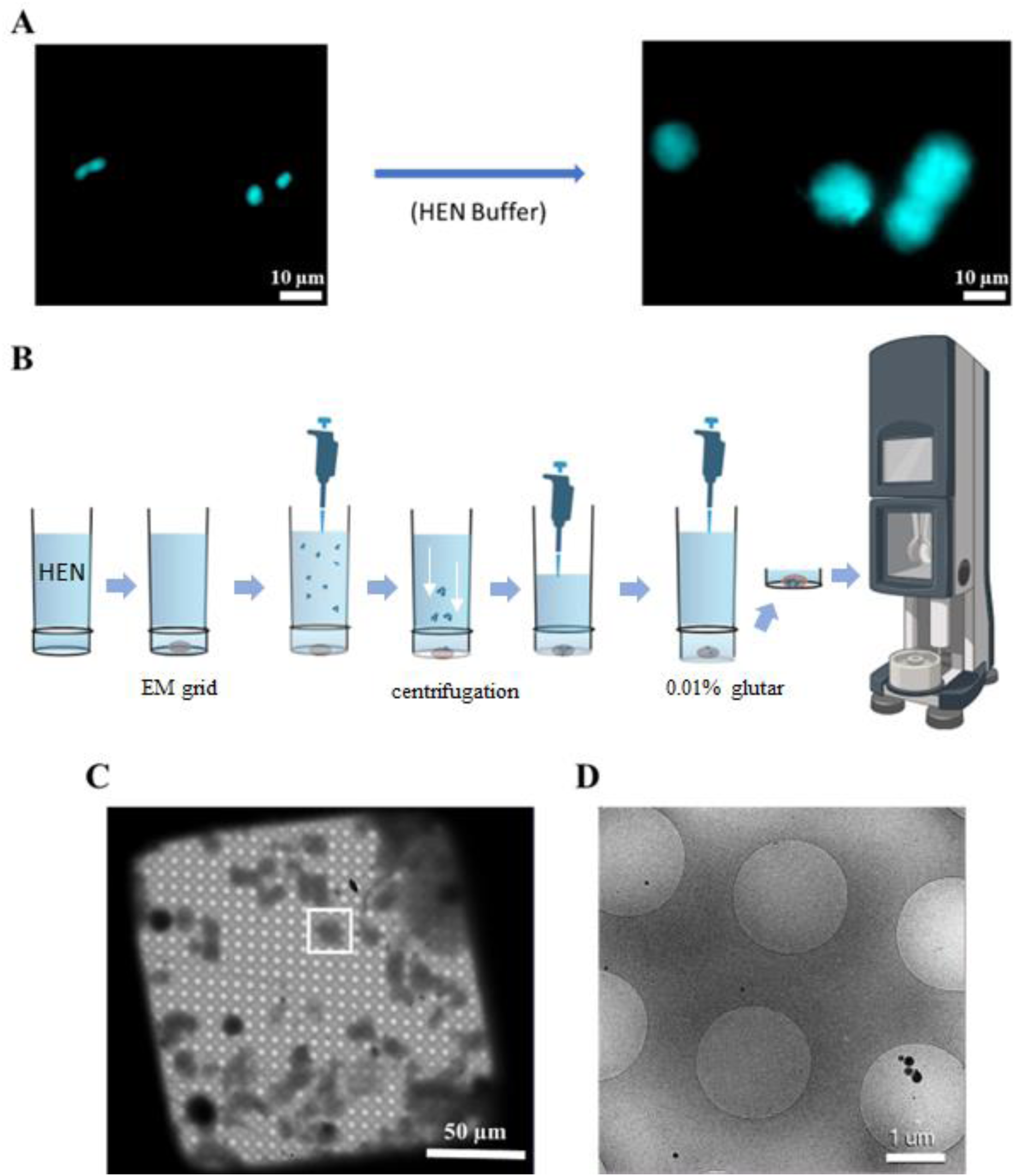
Chromosome preparation for cryo-EM. (A) Partial decondensation of mitotic chromosomes *in vitro* in a low ionic strength buffer. DT40 mitotic chromosomes in polyamine buffer (left) and after incubation in HEN buffer (right). DNA fluorescently labeled with Hoechst and shown in cyan color. The signal intensity of pdXS was increased to match one of the compact chromosomes in the polyamine storage buffer. (B) The preparation of pdXS for vitrification. The lid with the electron microscopy grid fits onto the cut Eppendorf tube (see Supplementary Figure 1 for details). The setup was filled with the HEN buffer, then polyamine chromosomes were added and incubated at 4℃ for 1 hour to get partially decondensed. The pdXS were spun down onto the surface of the cryo-EM grid. The half of the liquid was removed and replaced by HEN containing 0.02% of glutaraldehyde (glutar). After 30 min of fixation on ice, the bottom lid was removed, and the grid was picked up by the Vitrobot tweezers and immediately processed to vitrification. (C-D) a cryo-EM image of pdXS obtained with this pipeline. pdXSs can be recognized as round semi-transparent densities (D). The regions of pdXSs overlaid with holes in main carbon support were targeted for cryo-ET (see Figure 2).

The chromosome preparation strategy for cryo-EM analysis was built with a strategy to minimize the risk of damage and achieve a reproducible optimal density of chromosomes attached to the cryo-EM grid. We performed the partial decondensation in a homemade centrifugation container based on a 1.5 ml Eppendorf tube that was then directly used for depositing the chromosomes onto the cryo-EM grid (Figure 1B). This setup was adapted from (Kiseleva et al. 2007) where a similar technique was used to deposit yeast nuclei on the EM grids. The isolated chromosome sample, stored in a glycerol-containing buffer, was overlaid onto 400 µL of a HEN buffer inside the centrifugation container, inducing partial decondensation within the vial. The container was then placed in a centrifuge, spinning the partially decondensed chromosomes directly onto the cryo-EM grid positioned at the bottom cap. After centrifugation, the grid with attached chromosomes could be easily picked from the cap using tweezers and mounted into a plunge-freezing device, such as the Vitrobot Mark IV (FEI). By eliminating the need for pipetting, this approach minimized the risk of damage to the highly fragile decondensed chromosomes.

The increase in size during partial decondensation was considered when calibrating the dilution of original chromosome stocks. Overcrowding was prevented to allow proper removal of excess liquid during the blotting step of plunge freezing. Insufficient liquid removal can lead to poor vitrification and consequently suboptimal imaging quality. We found that chromosome stocks with an approximate concentration of 0.5 mg/ml (10 OD260) must be diluted 1:100 times in the HEN buffer to provide a reasonable density of chromosomes distributed in each square of the EM grid as shown in (Figure 1C). An example of an individual particularly decondensed chromosome (pdXS) is shown in (Figure 1D). Scattered individual pdXS could be identified as dark roundish spots when the grid is imaged at low magnification in an electron microscope.

Cryo-ET revealed that the thickness of the vitrified chromosomes within the tomograms ranged from 150-350 nm, with a maximum thickness at the centre of the pdXS that reduces towards the periphery. The chromatin occupied the complete volume of the vitrified layer and showed uniformly distributed intact nucleosomes (Figure 2A). The structural integrity of nucleosomes was confirmed by their direct observation after computational denoising with deep learning networks (Buchholz et al. 2019) that also enhanced the visualization of linker DNA. Nucleosomes could be recognized thanks to their characteristic different orientational views and their connection by the linker DNA (Figure 2B-I). Subtomogram averaging of nucleosomes identified in our tomograms using Template Learning (Harastani et al. 2024) and 3D template matching method yielded a subtomogram average resolution of ∼13 Å which is comparable to that obtained by other studies on native chromatin (Chen et al. 2024; Tan et al. 2023). Further classification of nucleosomes revealed expected conformational variability without obvious evidence of sample damage, supporting our qualitative assessment of the high quality of sample preparation.

**Figure 2:**
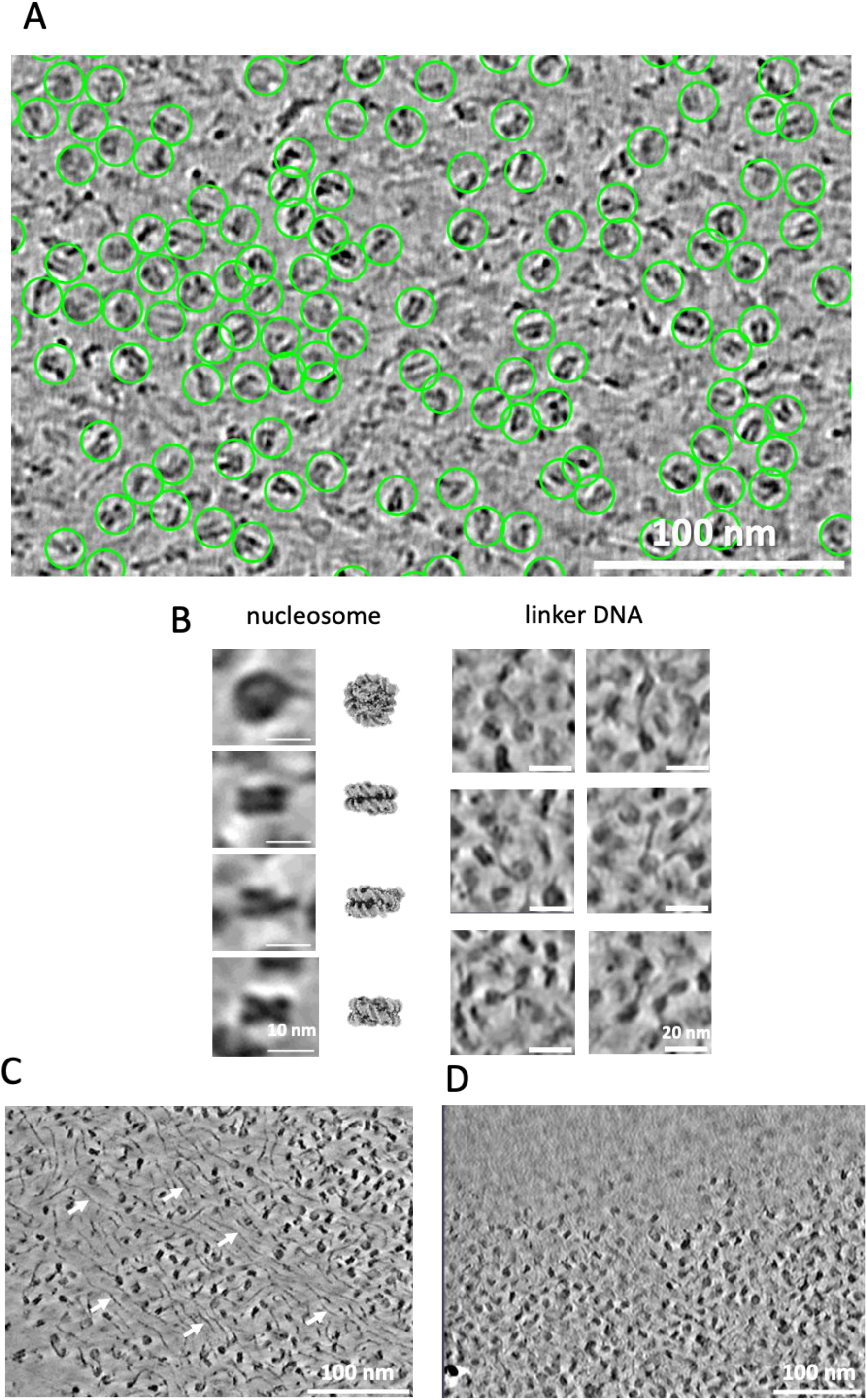
Visualization of chromatin integrity in pdXSr. (A) A 4.15 Å thick central slice of a denoised tomographic reconstruction. Green circles pointing at multiple nucleosomes automatically annotated with Template Learning software. (B) Characteristic views of individual nucleosomes and linker DNA extracted from (A) shown side by side with the corresponding orientation of the nucleosome atomic model (PDB 1EQZ). (C,D) Chromatin disassembly at the air-water interface resulting in long stretches of naked DNA visualized at the surface of the sample and are indicated by white arrows. (C) can be avoided by a gentle fixation of the sample with 0.01% glutaraldehyde before vitrification (D).

At the surface layers of chromatin exposed to the air-water interface, we observed long DNA stretches, indicating that some nucleosomes were destroyed due to the exposure to the air-water interface (Figure 2C). To minimise this damage, we employed a very gentle fixation using 0.01% glutaraldehyde. Such gentle fixation has been earlier used as a strategy to prepare samples for fragile macromolecular complexes while maintaining their structural details for single particle analysis (Papai et al. 2020). We noticed that gentle fixation with 0.01% glutaraldehyde (Kastner et al. 2008) performed after centrifugation (see Figure 1B) did not affect the chromosome decondensation, but eliminated the nucleosome denaturation artefacts at the sample surface (Figure 2D; see Materials and Methods for details).

### Fluorescent labelling of condensin proteins in pdXS shows axial staining

We used a chicken DT40 cell line containing Halo-tagged SMC2 condensin (Samejima et al. 2024) which could be fluorescently labeled using Halo TMR ligand. *In vivo* labelling of condensins with halo TMR ligand (Promega) in PAF-fixed nocodazole-arrested mitotic DT40 cells shows its axial localization along the two sister chromatids of the chromosomes (Figure 3AB). Our observations are in agreement with previously published results on the localization of condensins in mitotic cells using immunofluorescence staining and other fluorescent labelling approaches that have demonstrated that condensins are concentrated along the axial region of the mitotic chromosome in the mitotic cell (Maeshima, Eltsov, and Laemmli 2005a); (Hibino et al. 2024); (Walther et al. 2018); (Samejima et al. 2012)).

**Figure 3.**
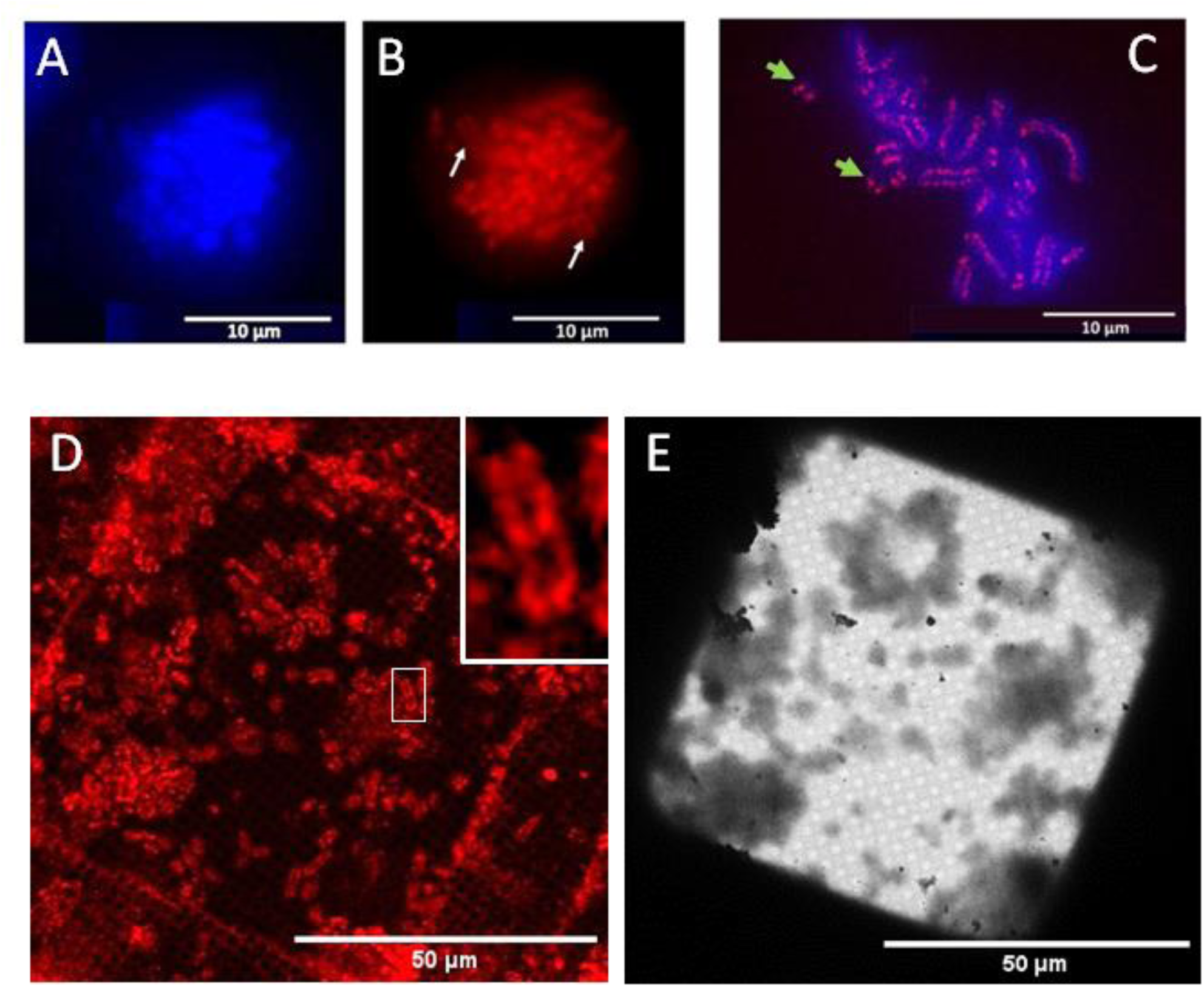
Fluorescence imaging and cryo-CLEM of SMC2-Halo-TMR. (A, B) Nocodazole-arrested DT40 cells stained in vivo with Halo-TMR and fixed with PFA. DNA counterstained with Hoechst is shown in blue (A). White arrows in (B) indicate examples of sister chromatids with linear axial staining. (C) Partially decondensed chromosomes (pdXS) stained during isolation and attached to coverslips show linear condensin axes staining. DNA is counterstained with Hoechst (blue). Green arrows highlight minichromosomes in a pdXS cluster. (D, E) Correlative light and electron microscopy (CLEM) of SMC2-Halo-TMR chromosomes attached to an EM grid. (D) Cryogenic imaging of a grid square using a Zeiss LSM900 Airy-scan confocal microscope axial staining of chromosomes. A zoomed-in view of a chromosome (outlined as a white frame in D) shows a characteristic X-shaped axial region (inset, white frame). (E) Corresponding cryo-EM image of the same grid square.

Our attempts at direct fluorescent staining of Halo-tagged SMC2 condensins with Halo TMR ligand during partial decondensation or directly on the EM grid for cryo-fluorescence imaging led to a very high background noise. Therefore, we established the halo-TMR fluorescent labelling of the SMC2 Halo-tagged condensin during the polyamine-based chromosome isolation procedure once the swollen cells were lysed by homogenization. The subsequent sucrose and percoll gradient purification steps removed the unbound stain. (Figure 3C) shows this labelling-during-purification in pdXS DT-40 chromosomes in HEN buffer. The continuous linear aspect of SMC2-Halo signal visible in intact cells remained in chromosomes partially decondensed in the HEN buffer.

### Cryo-ET of the axial regions reveals heterogeneous non-histone macromolecular densities

After labelling-during-purification, partial decondensation in HEN and gentle fixation, chromosomes were deposited on the surface of the EM grids, plunge frozen, and imaged with high-resolution confocal microscope (Zeiss LSM900 Airyscan 2) in cryogenic conditions. We found that the axial staining of SMC2-Halo-TMR was lost when copper grids were used. A uniform background staining was redistributed to the complete chromosome contours (Supplementary Figure 2A-D). We hypothesise that the loss of the specific SMC2-Halo-TMR staining in samples prepared on copper grids results from a reactivity of copper ions released by the grid with the HaloTag-ligand system. In contrast, on gold grids, the axial staining remained (Figure 3D). Cryo-fluorescence imaging showed two distinct strands of the axial signal corresponding to the two sister chromatids. Besides fluorescence, we collected refracted light images of the corresponding grid squares, which allowed us to use the position of regular holes in Quantifoil grids for localization and registration of the regions of interest in low magnification cryo-EM maps and to target regions for tilt series acquisition (Figure 3E, Figure 4A).

**Figure 4:**
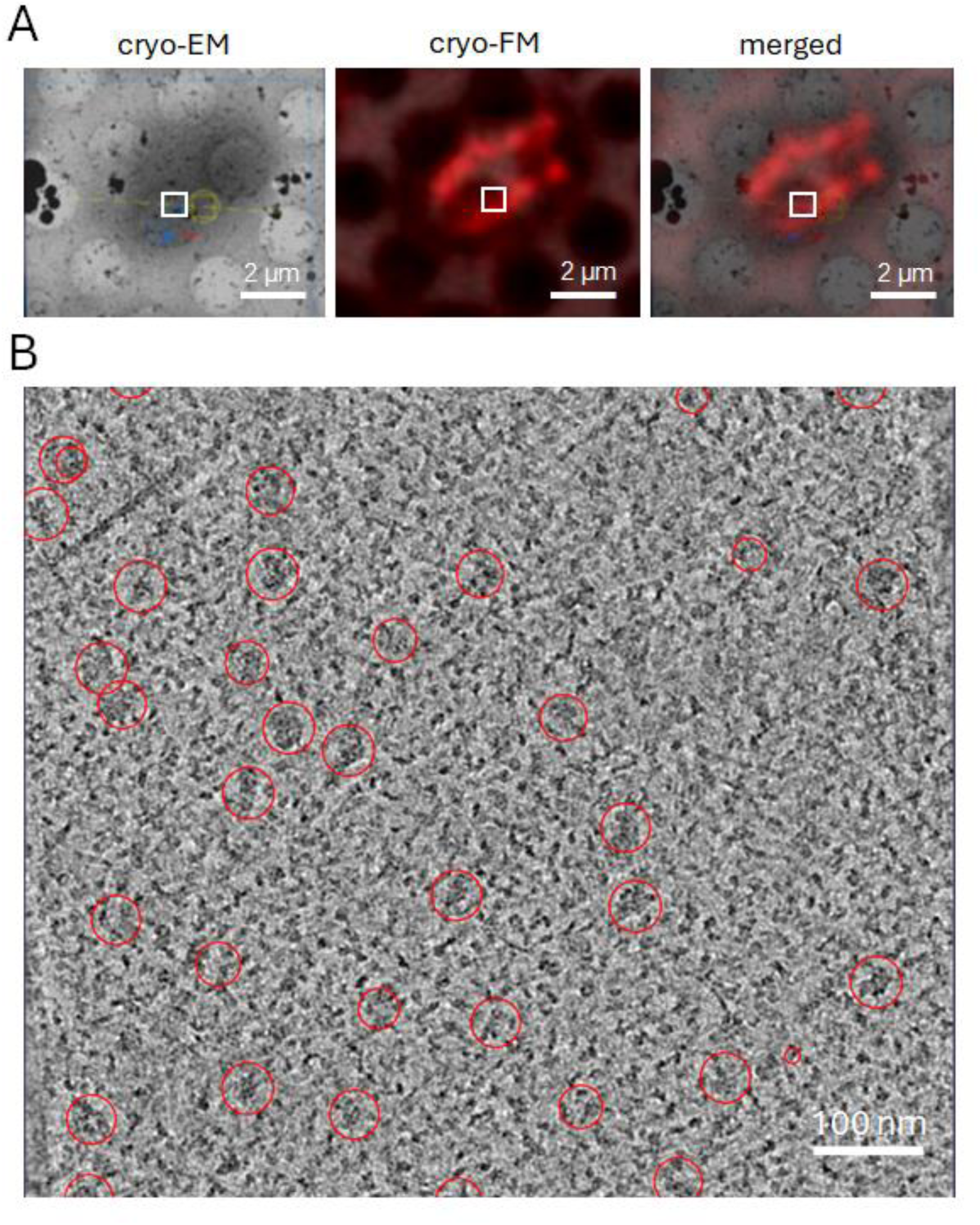
Cryo-CLEM of a condensin-rich axial region of pdXS shows heterogeneous non-histone macromolecular densities (NHMDs) in the bulk of chromosomes. (A) An overview image of the same vitrified pdXS imaged with cryo-EM at a low magnification (cryo-EM), with cryo-fluorescence confocal microscope (cryo-FM), and an aligned merged view are shown. The region of cryo-ET acquisition shown in (B) is indicated by a white frame. (B) A 4.15 Å thick central slice of a denoised tomogram acquired in the region indicated in (A). NHMDs are annotated in red spheres in a chromosomal environment. Interestingly, a majority of NHMDs are not interconnected, but spaced by chromatin regions. The position of this tomographic slice within the reconstructed volume is shown in Supplementary Figure 3A

Cryo-ET of the axial region of the chromosome showed abundant nucleosomes, linker DNA and larger densities that are distinguishable from nucleosomes, which are potentially part of the non-histone component of the mitotic chromosome (Figure 4B; Supplementary Figure 3A). These densities were large, measuring approximately 30–45 nm, interspersed with chromatin/DNA. We have termed them as Heterogeneous Non-Histone Macromolecular Densities (NHMDs).

The peripheral surface layers of chromosomes also demonstrated clusters of molecular densities of approximately 30 nm in size (Figure 5A; Supplementary Figure 3B). Such clustered densities had been reported by previous cryo-EM studies in isolated human chromosomes (Nishino et al. 2012). They were suggested to contain ribosome-related material that accumulates at the chromosome surface because they could be labelled with antibodies against the ribosomal component S6 antibody. Indeed mass spectroscopic analysis of isolated mitotic chromosomes shows the presence of ribosomal-related proteins (Ohta et al. 2010). Our chromosome preparation also showed S6-positive labelling (Figure 5B), supporting the presence of ribosome-related material at the chromosome periphery. Using fluorescence microscopy, the peripheral labelling did not show any correlation with the axial staining of SMC2-Halo-TMR, supporting the differential localization of these two sorts of densities in cryo-ET(Figure 5B).

**Figure 5.**
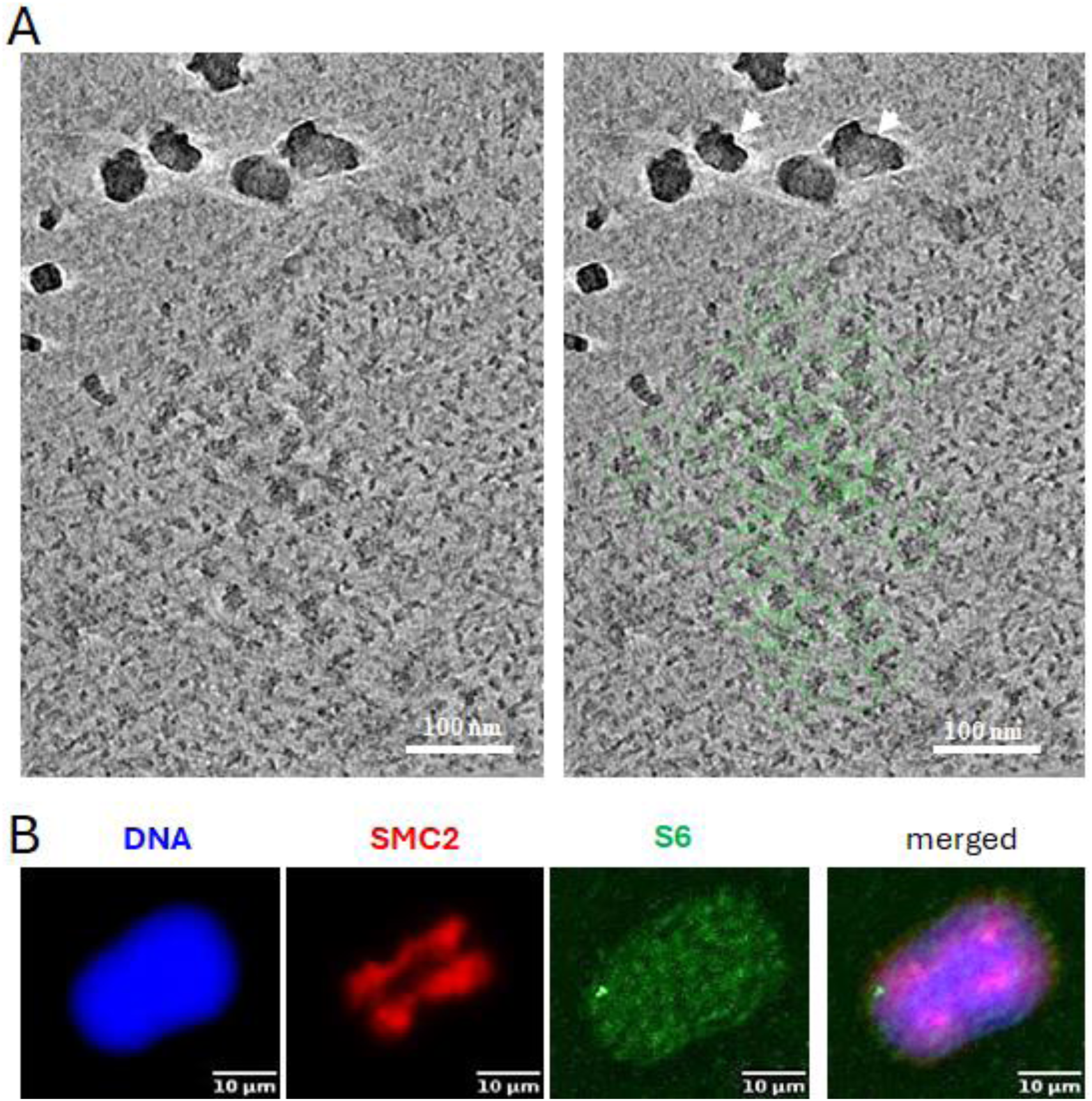
Ribosome-related densities aggregate at the chromosome surface. (A) A 4.15 Å thick tomographic slice through the surface of pdXS showing dense structures annotated as green circles on the right. Highly-contrasted dark densities as ice crystals accumulated at the surface of the sample during cryo-fluorescence imaging (white arrows). The position of this tomographic slice within the reconstructed volume is shown in Supplementary Figure 3B (B) Immunofluorescence labelling of pdXS with antibody S6 reveals clusters of ribosome-related material at the chromosome surface. DNA is counterstained with Hoechst, SMC2-Halo labelled with TMR during chromosome isolation.

Once we visually identified the presence of NHMDs within the bulk of pdXSs and the distinct clustered densities on their surfaces, we explored the possibility of their unbiased classification using quantitative tools to enable automated particle identification for subtomogram averaging. For this purpose, we manually annotated NHMDs and surface densities in three representative tomograms (TS_22, TS_19, TS_13) and combined them into a single group for each tomogram. Based on the observed clustering of densities near the chromosome surface, we explored two approaches: (a) analysis of a distribution of distances measured from the density and nearest chromosome surface; (b) clustering analysis of spatial distribution of densities using density-based clustering non-parametric algorithm (DBSCAN; (Shah 2012).

At first the distances from each density to the nearest surface were measured for each tomogram (Supplementary Figure 4). Interestingly, the distributions of TS_22 and TS_13 showed indications of bimodality, thus supporting the presence of two subpopulations of particles. This feature was less pronounced in TS_19, likely because the surface population of particles was lower. Since no statistical difference in distance distributions between tomograms was found using the “Kolmogorov–Smirnov” two-sample test, also merged all three distributions into one (Supplementary Table 2). The merged distribution also clearly indicated bimodality (Supplementary Figure 4, Merged). We applied a Gaussian Mixture Model (GMM) analysis (Y. Huang et al. 2005) is a probabilistic model used for interpretation to the distributions made of multiple Gaussian distributions. This allowed us to define two subpopulations of particles in relation to the nearest surface distance: the cumulative surface fraction (mean 23 nm, SD 14 nm, Figure 6A, NSD) appears more “focused”, whereas the inner fraction of particles (mean 117 nm, SD 38 nm) are dispersed with a relatively random “flat” distribution. Applied independently, DBSCAN also identified large clusters of closely located particles in the vicinity of the chromosome surface (Figure 6A DBSCAN, Supplementary Figure 5), whereas no meaningful clustering was revealed inside the volume of the pdXS. Interestingly, densities assigned into the surface fraction by the first method and the combined surface clusters identified by DBSCAN showed up to 93.5% of overlap (Figure 6A NSD+DBSCAN). A side-by-side comparison of images of the densities extracted from the NHMDs fraction and the surface fraction demonstrates distinct structural differences (Figure 6B): NHMDs display greater heterogeneity in shape and are generally larger than particles in the surface fraction. Some of the particles extracted from the surface fractions also exhibit morphological similarities to ribosomes, supporting the possibility that these are ribosome-related clusters. These results support the feasibility of quantitative separation of NHMDs from surface components.

**Figure 6:**
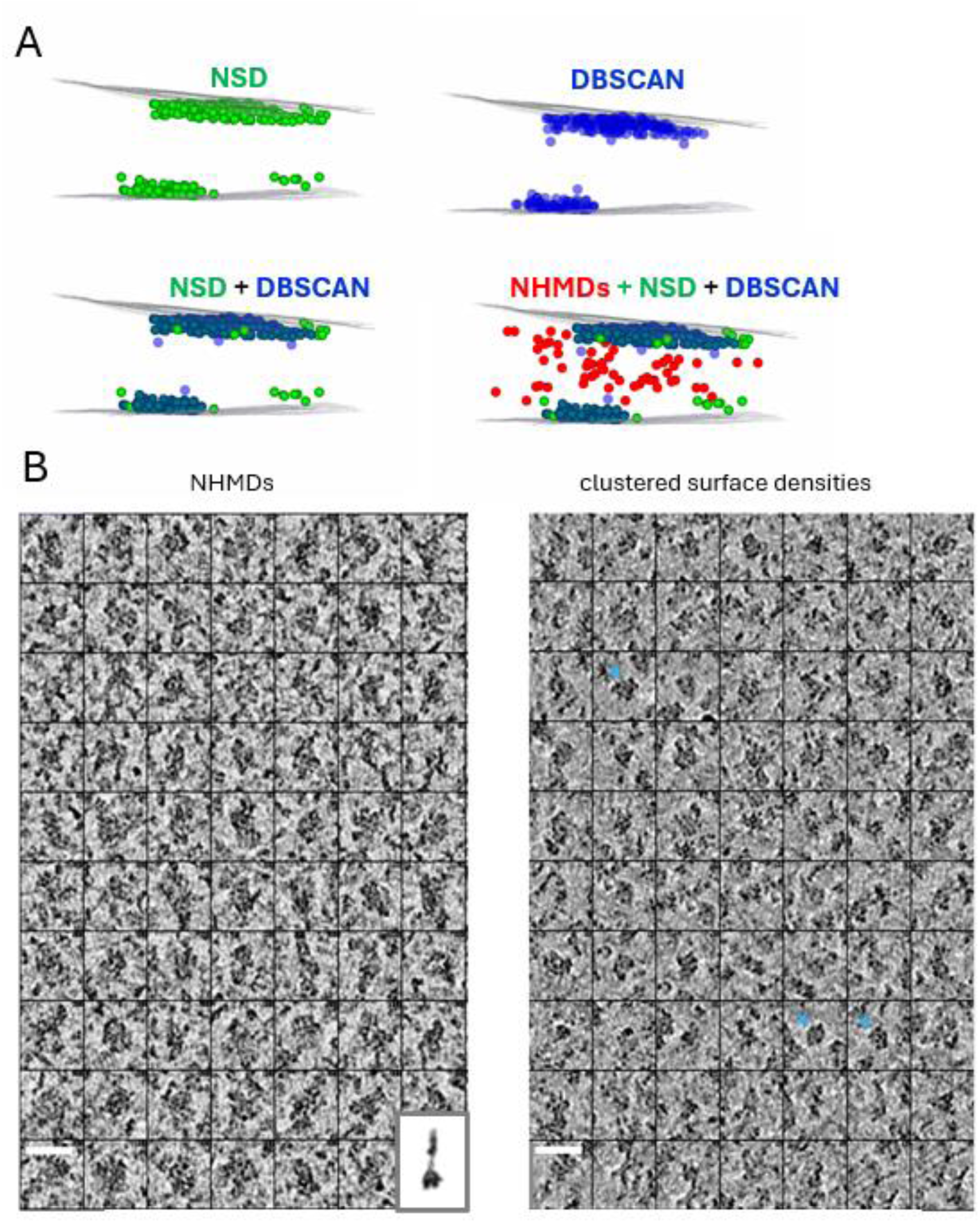
Discrimination of NHMDs and ribosome-related surface densities based on their spatial distribution and clustering behaviour. (A) Densities of the tomogram TS_22 were classified to the surface fraction based on the analysis of their nearest surface distance (NSD, green spheres) and clustering analysis (DBSCAN, blue spheres) showing 93.5% of overlap (dark-green spheres). The surface fractions obtained by both methods were excluded and remaining densities were assigned to the NHMDs fraction (red spheres) which is dispersed inside the chromosome volume. The surfaces of the chromosomes are shown in grey (see Material and Methods for details). (B) Galleries of 4.15 Å thick central slices of subtomograms randomly selected NHMDs and clustered surface fraction extracted from TS_19. NHMDs appear to be generally larger and more pleomorphic than the clustered surface densities. Notice that NHMDs are also larger than yeast condensin complex (Lee et al. 2020, PDB 6YVU) shown in the corresponding scale as an inset in the gallery. Blue arrows indicate some dense compact particles that are often found in the surface fraction. Scale bars in both galleries are 30 nm.

Our attempts to apply subtomogram averaging of 1000 NHMDs did not converge to any interpretable structure, further supporting their high structural heterogeneity. Visually, we did not observe any other components except the NHMDs, nucleosomes and linker DNA in the bulk of the axial region of the mitotic chromosomes.

NHMDs were separated by regions of chromatin. The average nearest neighbour distances between the centres of NHMDs calculated for three representative tomograms TS_22, TS_19 and TS_13 with the best alignment scores (<1 nm) are 70 nm, 62 nm and 98 nm respectively. Considering the 35–40 nm size of NHMDs, these results indicate that NHMDs are separated by approximately 40 nm on average (Supplementary Figure 6).

While NHMDs were abundant in the axial areas, control tomograms recorded outside of the axial areas show only a few potential densities (data not shown). This observation together with the fact that no other non-nucleosomal macromolecular complexes were distinguished in the bulk of chromosomes suggests that at least a part of NHMDs could be related to condensin complexes, although DNA topoisomerase II and KIF4A could also contribute to these densities.

## Discussion

We have adapted the polyamine-based chromosome isolation method of Lewis and Laemmli (Lewis and Laemmli 1982) and have developed a pipeline for the deposition of partially decondensed chromosomes (pdXS) on a cryo-EM grid for high-resolution visualization of the internal structure of chicken DT40 mitotic chromosomes by cryo-ET. Structural preservation at the molecular scale was confirmed by the integrity of nucleosomes, which are relatively sensitive to mechanical damage (Chien and van Noort 2009). The preservation of nucleosome structure in our preparations was validated by subtomogram averaging, which achieved an approximate resolution of 12.8 Å, with local resolutions ranging from 12 to 20 Å (Harastani et al. 2024). Furthermore, by using gentle GraFix crosslinking (Kastner et al. 2008) we eliminated the chromatin disruption at the air-water interface, visible in previous studies (Beel et al. 2021).

Partial chromosome decondensation enabled us to distinguish non-histone components within the central axial regions of pdXS. Indeed, we could only identify three components in the internal regions of our chromosome preparation: nucleosomes, linker DNA, and additional molecular densities that showed extremely high structural heterogeneity. We have termed them heterogeneous non-histone molecular densities (NHMDs). We hypothesize that they constitute the non-histone component of mitotic chromosomes. These NHMDs have not been previously described in *in situ* cryo-ET studies of mammalian mitotic chromosomes (Chen et al. 2024). This may be attributed to the high chromatin crowding in mammalian chromosomes, which could prevent the identification of such complexes. Consistent with this hypothesis, an *in situ* cryo-ET study reported multimegadalton globular complexes within mitotic chromosomes of *Schizosaccharomyces pombe* (Cai et al. 2018).

However, this explanation does not apply to similar work on human chromosomes (Beel et al. 2021), where a similar isolation method using polyamines and partial chromosome decondensation was employed. Based on previously published data (Beel et al. 2021), the degree of chromatin compaction appears comparable to that observed in our work. In this work, there is no indication of the localisation of the data collection relative to the chromosome axis, therefore the lack of additional molecular densities seen may be due to the data having been collected from the thinner peripheral regions of the chromosomes. Human chromosomes are significantly larger (with the smallest and the largest chromosomes being ∼ 46 Mb and ∼ 250 Mb) (Nurk et al. 2022) than most of the chicken chromosomes. Chicken cells have numerous microchromosomes (with an average size of ∼10.2 Mb) and dot chromosomes (with an average size of 4 Mb) (Z. Huang et al. 2023). Their small size should allow multiple partially decondensed chromosomes to fit within a single EM grid square without overlapping (Figure 1C), making tomographic data collection more efficient. Thus the chicken chromosomes offer an opportunity for cryo-ET analysis of whole chromosomes without the need for additional sample thinning, by cryo-FIB-SEM or cryo-sectioning.

We noticed that NHMDs have lower contrast in cryo-EM tomograms in comparison to nucleosomes. The latter contains DNA that produces higher scattering because of phosphorus atoms. NHMDs therefore are more difficult to distinguish in crowded chromosomes, possibly providing another reason for NHMDs having not been previously identified in *in situ* tomograms (Chen et al. 2024). On the other hand, this observation strongly suggests that NHMDs are densities formed from protein alone. Combined with the presence of fluorescent signals from Halo-SMC2, this observation allows us to hypothesize that condensin proteins contribute to the formation of at least some of these densities. In our study, we could not distinguish condensin I and II, as SMC2 is found in both. However, considering that condensin I is 5 times more abundant than condensin II (Walther et al. 2018), we could preferentially look at condensin I, as a constituent part of some of the NHMDs.

The dimensions of NHMDs are substantially larger than the available structures of condensin. Also, the structure of the chicken condensin complexes are not known, yeast condensin is smaller (∼15 nm in the clamped state) (Shaltiel et al. 2022) than NHMDs, suggesting a possibility of multimerization. Previously published data has shown evidence for the existence of multimerization in condensins purified from yeasts (Keenholtz et al. 2017). Alternatively, or in addition, the NHMDs could consist of other non-histone proteins associated with condensin. Condensin I and the chromokinesin KIF4 are known to interact with one another. It is also possible that they act with DNA topoisomerase IIα to shape the mitotic chromosome (Samejima et al. 2012).

The presence of ribosome-associated densities on the surface of chromosomes complicates the observation of structurally relevant non-histone proteins in mitotic chromosomes. However, we developed a computational framework that enables their distinction from NHMDs, which will facilitate automated processing of NHMDs in the future.

Our observations of discrete and spaced NHMDs does not support the presence of a continuous scaffolding protein structure in mitotic chromosomes (J. R. Paulson and Laemmli 1977), although some rearrangement of the axial structure during partial decondensation cannot be excluded. Indeed, an earlier study has used immuno-gold electron microscopy on pdXS to show that the scaffold component DNA topoisomerase II formed a diffuse network of discrete complexes (Earnshaw and Heck 1985). Subsequently, (Poirier and Marko 2002), (Samejima et al. 2024) (Walther et al. 2018) proposed that no continuous proteinaceous structure is present in the axial region of the chromosome as part of the mitotic chromosome scaffold. We suggest that the overall chromosome organization is largely preserved during decondensation is based on two observations: (i) partial decondensation of chromosomes in HEN buffer maintained the linear SMC2-Halo-TMR signal visible in conventional fluorescence microscopy; and (ii) partial decondensation of isolated mitotic chromosomes is a reversible process, as shown in previous studies (Earnshaw and Laemmli 1983), (Beel et al. 2021).

This study represents the first cryo-EM visualization of non-histone complexes directly within mitotic chromosomes and establishes a sample preparation pipeline for their future studies. Our findings open a way towards the study of the internal structure of relatively unexplored axial regions of pdXS using cryo-CLEM. We believe that the advancements in image processing software will enable us to address the heterogeneous structures of NHMDs and resolve their molecular organization in the axial region of the mitotic chromosome.

## Materials and Methods

### Cell Culture and synchronizing cells to mitosis

DT40 cells were cultured at 39°C in a 5% CO2 atmosphere in Roswell Park Memorial Institute (RPMI) 1640 medium supplemented with 10% (v/v) fetal bovine serum (FBS), 1% chicken serum and 1 % Penicillin: Streptomycin in a humidified environment (Ohta et al. 2010). The cell density is maintained between 1-10×10^5^ cells /ml to maintain the cells in a log phase. DT40 cells were synchronized using 0.5 μg/ml of nocodazole for 12-14 hours. This strategy gives a mitotic index of > 60%.

### Mitotic chromosome isolation

Mitotic chromosome isolation was done based on the polyamine-based purification method (Lewis and Laemmli 1982). The synchronized cells at a cell density of 5×10^8^ were harvested by centrifuging the culture at 1600 g for 5 mins at 4°C. The cells were swollen at room temperature for 5 minutes in 45 ml of low salt buffer containing 7.5 mM Tris:HCl pH 7.4, 40 mM KCl, 1 mM K-EDTA pH 7.4, 0.375 mM Spermidine, 0.15 mM Spermine followed by pelleting of the swollen cells at 900 g for 3 minutes at 4℃. The swollen cells are resuspended in a lysis buffer containing 15 mM Tris:HCl pH 7.4, 80 mM KCl, 2 mM K-EDTA pH 7.4, 0.75 mM Spermidine, 0.3 mM Spermine and 0.1% NP40 followed by lysis using a dounce homogenizer. The nuclei from the lysed non-mitotic cells were separated by centrifugation at 200 g for 5 minutes at 4℃. The supernatant was overlaid on a step sucrose gradient (15%, 60%, and 80% [w/v]) in a buffer containing 5 mM Tris:HCl pH 7.4, 2 mM KCl, 2 mM K-EDTA pH 7.4, 0.375 mM Spermidine and 0.1% NP40 and subjected to sucrose gradient centrifugation ag 1600 g for 30 minutes. The 60-80% interface was recovered and was mixed with a self-forming percoll gradient in a buffer containing 5 mM Tris:HCl pH 7.4, 2 mM KCl, 2 mM K-EDTA pH 7.4, 1.855 mM Spermidine, 0.592 mM Spermine, 0.1 % NP40 made in Percoll solution (from Cytiva, Catalogue number-17-0891-01). The sample was centrifuged at 44,000 g at 4℃ for 45 minutes. The band containing the chromosomes was recovered and washed in a buffer containing 5 mM Tris:HCl pH 7.4, 2 mM kcL, 2 mM K-EDTA pH 7.4, 0.375 mM Spermidine and 0.1% NP40 to remove the excess of percoll. The isolated chromosomes are stored in a buffer containing 3.75 mM Tris:HCl pH 7.5, 20 mM KCl, 0.5 mM EDTA, 0.05 mM Spermine, 0.125 mM spermidine, 0.1% NP40 and were stored in 60% glycerol at −20℃.

### Chromosome cryo-EM sample preparation

Quantifoil 200 mesh gold grids (R 2/2; Quantifoil Micro Tools GmbH, Germany) covered with 2 nm thin continuous carbon film were used for chromosome preparations and were glow-discharged immediately before use in a gas mixture of 80% Argon:20% Oxygen.

Partial decondensation of isolated mitotic chromosomes was performed by 100-fold dilution in HEN (10 mM Hepes pH 7.5, 0.2 nM EDTA, 25 mM NaCl) buffer containing 0.1% NP 40 and 0.5 mM ATP ƔS. After 1 hour incubation at 4 °C, mitotic chromosomes were spun down onto the surface of the glow-discharged Quantifoil 200 mesh copper and gold grids (Quantifoil Micro Tools GmbH, Germany) at 720 g for 5 minutes. The mild fixation with glutaraldehyde (EMS) was added to the solution at a final concentration of 0.01% in the same buffer with 0.1% NP40 and was carried out on ice for 30 mins. 10 nm BSA Gold Tracer (EMS, Hatfield, PA, USA) was added to the grid at a 5:1 ratio (chromosome:gold). The grid was blotted from the carbon side using a teflon sheet and the metal side using a blotting paper respectively similar to as shown in (Schaffer et al. 2015). The grid was blotted for 25 seconds with a blot force of 10 and flash-frozen into liquid ethane using Vitrobot Mark IV (Thermo Fisher Scientific) at 4 °C and 100 % humidity. The same preparation was used for chromosomes stained with Halo-TMR ligand and unstained.

In our initial cryo-EM experiments we used Multihole Quantifoil 200 mesh copper grids (R 2/2; Quantifoil Micro Tools GmbH, Germany). We found that specific axial SMC2-Halo-TMR staining was lost when copper grids were used while nucleosome structure was not affected.

### Cryo-ET data acquisition and processing

The tilt series was recorded on Titan Krios G4 (Thermo Fisher Scientific) with Falcon 4 camera equipped with a Selectris X energy filter, Titan Krios G3 (Thermo Fisher Scientific) equipped with a Quantum energy filter and Gatan K3 detector using SerialEM Version 4.1.0 beta 58. The images were acquired at a pixel size of 2.075 Å/pixel at −2 to −4.0 μm defocus at a dose rate of 3.2 e−/Å2 per tilt image fractionated over 10 frames. A dose symmetric tilt scheme (Hagen, Wan, and Briggs 2017) was used with a 3-degree increment step and the tilt range was set to ±60 degree using Serial EM (Mastronarde 2005).

The movies are motion-corrected and averaged to make a stack using MotionCor2 (Zheng et al. 2017). The tilt-series are aligned and reconstructed in Etomo using weighted back projection with SIRT-like filters applied (Kremer, Mastronarde, and McIntosh 1996). The reconstructions were denoised with cryoCare (Buchholz et al. 2019). Nucleosomes were picked using Template Learning and 3D template matching (Hrabe et al. 2012), (Best, Nickell, and Baumeister 2007), (Harastani et al. 2024) and subtomogram averaging was performed in Relion 4 (Kimanius et al. 2021).

### Fluorescence labelling of Halo-SMC in mitotic cells and isolated mitotic chromosomes

For the fluorescent labelling of condensins in the chicken DT40 SMC2-Halo (Samejima et al. 2024) nocodazole-arrested mitotic cells, 1:10,000 dilution of 5 mM Halo-TMR ligand (from Promega, Catalog Number: G8252) was added to nocodazole-arrested cell culture. The cells were incubated with the Halo-TMR ligand for 30 mins while in the culture *(in vivo* labelling). The excess of the halo-TMR dye present in the culture media was removed by harvesting the cells by centrifugation at 1800 g at 39°C for 15 minutes and replacing the supernatant media with fresh prewarmed media. These steps were followed 3 times to completely remove the Halo-TMR ligand from the media. The cells were finally harvested by centrifugation but resuspended in PBS at room temperature. The cells were spun down to the surface of poly-l-lysine coated coverslip using 2 ml buckets followed by fixation with 0.8% formaldehyde in PBS at room temperature. DNA was fluorescently stained with 1 μg/ml of Hoechst 33258 (Sigma-Aldrich) in PBS for 10 minutes at room temperature. The coverslips were mounted to a 35 mm petri dish with a 10 mm hole.

For fluorescent labelling of the isolated mitotic chromosomes, the staining was conducted during the chromosome isolation steps so that the extra dyes could be subsequently removed during the chromosome purification steps. Once the cells are lysed using a dounce homogenizer, 1:20,000 dilution of the 5 mM of Halo-TMR ligand was added to the sample and incubated for 1 hour at 4℃. The standard isolation procedure (see Mitotic chromosome isolation) was then resumed from the sucrose gradient centrifugation. For preparing the chromosome for imaging, the chromosomes are partially decondensed in TEEN or HEN buffer in the presence of 0.1% NP 40, 0.5 mM ATP ɣS 4 ℃ for 1 hours and are spun down on the surface of poly-L-lysine coated coverslips (from EMS, Catalogue number: 72292-01) at 720g for 5 minutes using 2 ml buckets followed by fixation with 0.8% formaldehyde. The DNA was fluorescently stained with 1 μg/ml of Hoechst 33342 (from Sigma) for 10 minutes at room temperature. The coverslips were mounted to 35 mm Petri dishes with a 10 mm hole.

### Immunofluorescence labelling

The immunofluorescence labelling protocol was adapted from (Maeshima, Eltsov, and Laemmli 2005b, 2005a). Isolated DT40 chromosomes were partially decondensed in HEN buffer in the presence of 0.1% NP40 and 0.5 mM ATP Ɣs at 4 ℃ for 1 hours, then spun down on the surface of poly-L-lysine coated coverslips at 720 g for 5 minutes. The samples were fixed using 0.8% formaldehyde (EMS) in the HEN buffer at room temperature for 15 minutes. Then this fixation buffer was carefully removed from the top of the coverslips without letting it gry and 250 μl of XBE2 buffer (10 mM HEPES–KOH (pH 7.7), 100 mM KCl, 5 mM EGTA) containing 0.5 mM ATPγS and 1mM glycine was overlaid on the coverslip for 5 minutes at room temperature followed by washing with XBE2 buffer. The chromosome sample is blocked using 3% NGS (ThermoFisher Scientific, Catalogue number: 50197Z) in the XBE2 buffer at room temperature for 30 minutes. The coverslips were then subsequently incubated with an Alexa Fluor 488-conjugated Ribosomal Protein S6 antibody (Santa Cruz, Catalog No. sc-74459) was used at a 1:10,000 dilution. This step was followed by extensive washing with XBE2 buffer (six times, 3 mins each), the DNA was stained with 1 μg/ml of Hoechst for 10 minutes at room temperature.

### Optical microscopy at room temperature

The coverslips with labelled cells or isolated chromosomes were mounted to 35 mm Petri dishes having the central 10 mm hole. The dishes were filled with the corresponding buffer (XBE2, TEEN or HEN) to avoid sample drying. The samples were imaged with a Zeiss AxioObserver microscope. Z-stacks were acquired at a pixel size of 124 nm and a z-step size of 400 nm for TMR fluorescence and transmission light illumination. The images were deconvoluted using the inbuilt deconvolution tool from Zeiss. Fiji (Schindelin et al. 2012) software was used for preparing the maximum intensity projection of the stacks and for preparing the images.

### Cryo-fluorescence microscopy and image processing

Cryo-fluorescence data acquisition was performed on a Zeiss LSM900 Airyscan 2 microscope equipped with a Linkam cryostage CMS 196 and a 100× objective (numerical aperture 0.75) with an Airyscan module. The grid squares with optimally distributed chromosomes were selected, Z-stacks were acquired at a pixel size of 124 nm and a z-step size of 400 nm for TMR fluorescence and reflective light illumination, and processed with the Airyscan processing module within Zen software. The frame alignment of the image stacks was refined using the “Sift” function of Fiji (Schindelin et al. 2012) (Lowe 2004) and the maximum intensity projections of the stacks were made with Fiji and used for the localization of regions of interest in cryo-EM. The TMR and transmission light illumination channels were merged to relate the positions of chromosome axial regions with regular patterns of holes in carbon film of Quantifoil grids. It allowed identification of the regions of interest in low magnification cryo-EM maps of previously defined grid squares. Then the position of tilt series was defined and tilt series were collected using Serial EM and processed exactly as described above (*Cryo-ET data acquisition and processing)*.

GIMP (software version 2.10) was used to overlap the square maps obtained from the cryo-CLEM to the same square maps acquired with the cryo-electron microscope. The grid bars and the holes on the carbon were used as fiducials for alignment.

### Classification of non-nucleosomal densities

3 tomograms with the best alignment score < 1 nm (TS_22, TS_19 and TS_13) were selected. All the visually observed non-histone densities that could contain both NHMDs and ribosome-related densities were manually picked in the denoised volumes in IMOD. XYZ coordinates of all particle centres were manually picked in each tomogram and saved in the same model file in IMOD to have an unbiased classification.

#### Density-to-the-nearest surface distance analysis

Both the upper and lower chromosome surfaces were segmented manually in the slicer window of IMOD on X-Z planes of the tomograms. Distances from each picked density to both chromosome surfaces were measured using the mtk command of IMOD to find the nearest distance from the surface, and the minimum distance was selected for further analysis.

Histograms of the distances were calculated and plotted using custom Python script. Skewness and kurtosis metrics were computed to assess the asymmetry and tailedness of the distributions using Statistical Moments Calculation (Supplementary Table 1). Comparisons of individual tomogram distributions were performed using the Kolmogorov–Smirnov two-sample test, with the null hypothesis - “the two experimental distributions originate from the same underlying distribution”. As the null hypothesis could not be rejected based on the obtained P-values (Supplementary Table 2), the three distributions were merged for combined analysis.

Gaussian Mixture Model (GMM) (Y. Huang et al. 2005) was applied to separate two components of the bimodal distributions that were obtained. When applied to TS_22, it resulted in a threshold of 57 nm. For TS_13 and the combined distribution, GMM gave 42.44 nm and 45.43 nm (Supplementary Figure 4). The GMM threshold calculated for TS_19 was not considered, because of the absence of obvious signs of bimodality. Each of the distributions has been segregated into two fractions: a surface fraction (that could contain the ribosomal-related densities) and an inner fraction (that could contain the NHMDs) of particles using the largest threshold of 57 nm. Based on a given threshold (57 nm), the particles were blindly assigned to two indexes indicating whether they belonged to the surface or internal fraction in the axial region of the pdXS (Figure 6A, Supplementary Figure 4).

#### DBSCAN clustering

The DBSCAN clustering algorithm was selected due to its ability to identify clusters without predefined class numbers, minimizing potential bias (Deng 2020). All the visually observed non-histone densities were manually annotated that could contain the NHMDs and the ribosome-related densities were combined into a single point cloud for analysis. The optimal ɛ value was determined using the k-nearest neighbour (k-NN) distance plot, identifying a characteristic "elbow" point marking the transition from densely packed to more dispersed regions (Supplementary Figure 5A). This ɛ value was used to balance clustering sensitivity while avoiding over-or under-segmentation. The MinPts parameter, representing the minimum number of points required to form a cluster, was set to 2, to maximize the sensitivity of the cluster detection. Applied to TS_22 (ɛ =114 pixels) DBSCAN produced 57 clusters, between those the 3 largest clusters contained 128, 64 and 27 points, while other classes contained 2-3 points (Supplementary Figure 5B). Backmapping of the large classes revealed their surface localization. The inner volume showed weak clustering behaviour (Figure 6A). We separated the large classes by arbitrary thresholding and combined them in a single fraction 1, while combining all remaining to fraction 2. Similar clustering patterns were observed in tomograms TS_19 and TS_13. Then the overlapping ratio with the results of Density-to-the nearest surface distance analysis were calculated.

## Author Contribution

Conceptualization: ME, WCE, GP Methodology: ME, MH, KS, LR, WCE

Sample preparation and data acquisition: GP, MH, ME, CC, AD, NM, IL, AS, ZhY, WH

Results interpretation and validation: GP, MH, NTC

Funding acquisition: ME, WCE

Supervision: ME

Writing – original draft: GP

Writing – review & editing: GP, MH, LR, KS, NTC, WCE, ME

## Acknowledgments

The work was supported by ANR (ANR-20-CE11-0020-02 to MH and ME, ANR-23-CE45-0012-01 to ME), LabEx (ANR-10-LABEX-30-ME to GP and ME), the French Infrastructure for Integrated Structural Biology (FRISBI ANR-10-INSB-005) and Instruct-ERIC; and iNEXT-Discovery PID22849, grant number 871037, funded by the Horizon 2020 program of the European Commission. KS, LR, WCE were supported by Wellcome grants 107022 and 221044 to WCE and 203149 to the Wellcome Centre for Cell Biology. We acknowledge the access and services provided by the Imaging Centre at the European Molecular Biology Laboratory (EMBL IC), generously supported by the Boehringer Ingelheim.

