## Supplemental Figures and Tables for "Organisation of axial regions of isolated mitotic chromosomes visualised by cryo correlative light and electron tomography"

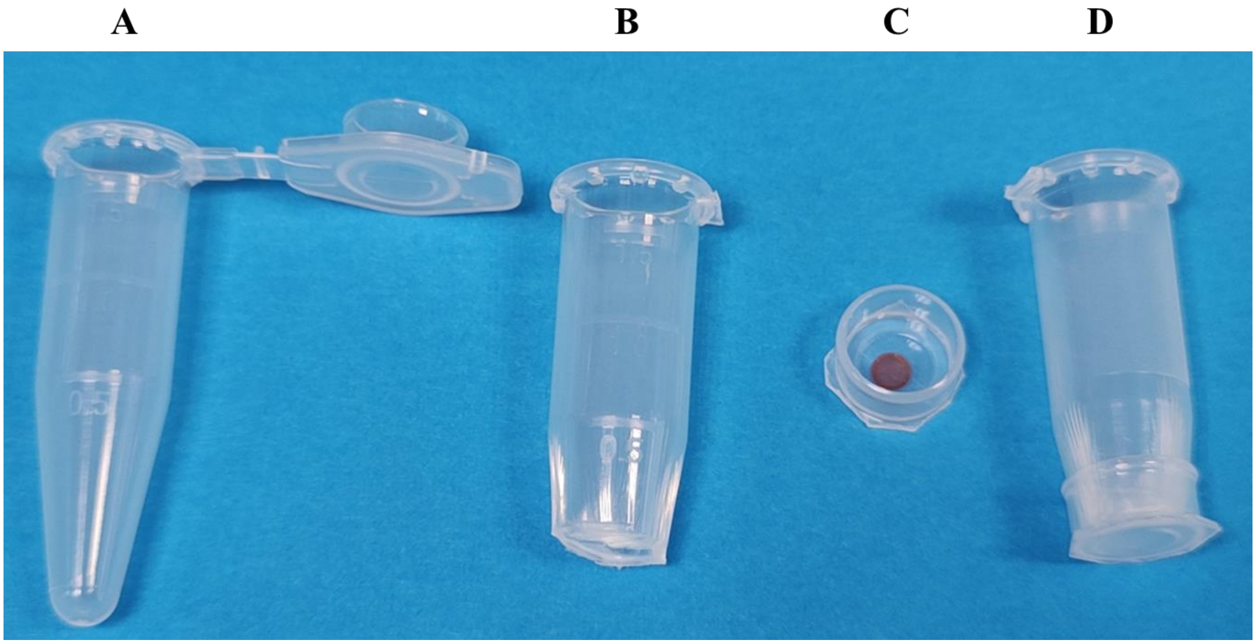


**Supplementary Figure 1.** Constructing a microtube chamber for spinning down chromosomes onto the surface of an electron microscopy grid for plunge freezing. (A) Removal of the lid and bottom part from a 1.5 ml Eppendorf microtube. The remaining lid attachment is cut off from the top of the tube. (B) The bottom of the tube is cut precisely to a diameter that could allow the lid to push in tightly to avoid leakage during centrifugation. (C) The electron microscopy grid is placed on the lid. (D) The lid with the electron microscopy grid fits onto the cut Eppendorf tube.

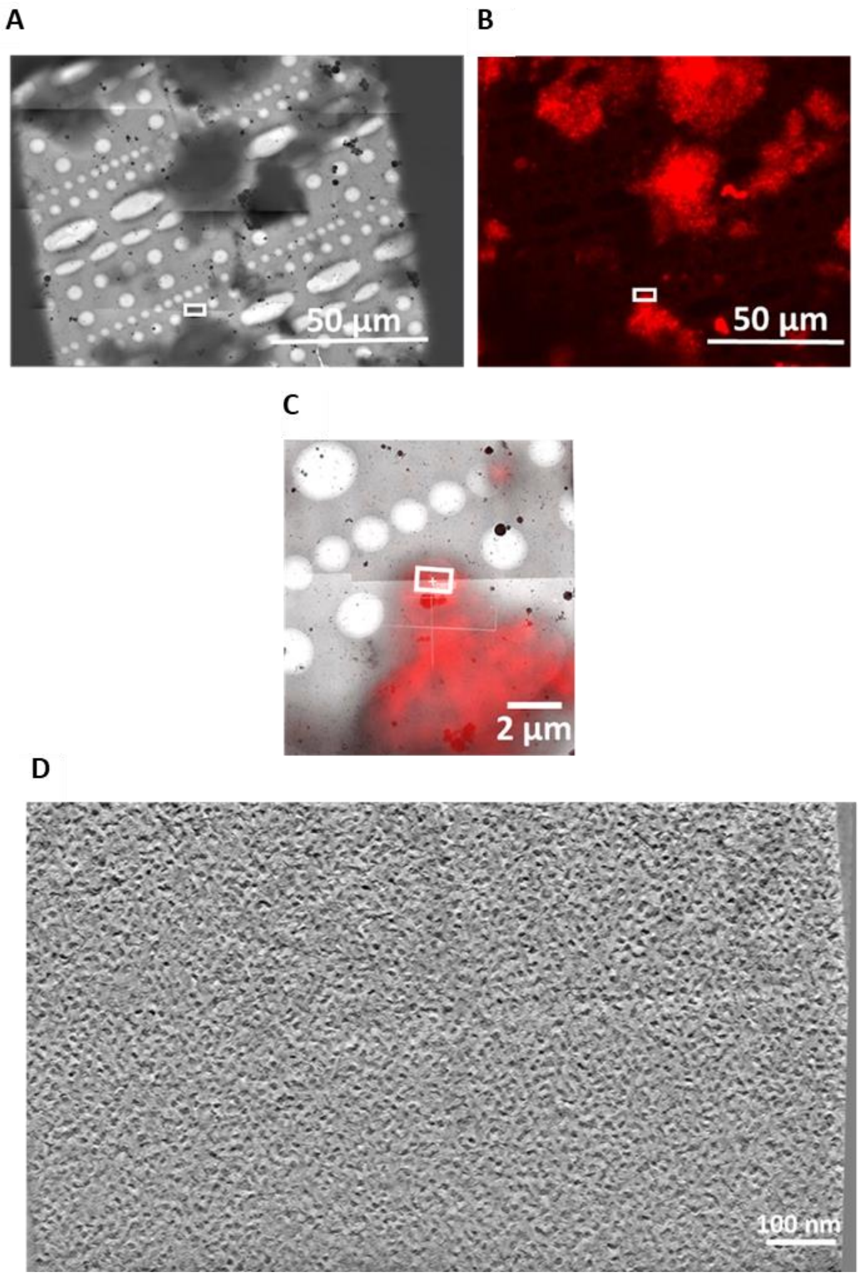


**Supplementary Figure 2.** (A) SMC2-Halo-TMR staining in chromosomes partially decondensed vitrified on cryo-EM grid with copper support. (A-C) Loss of the specificity of Halo ligand TMR staining of the axial regions of chromosomes after the partial decondensation and spinning down to EM grids with copper support. Interestingly, despite the loss of axial staining, the chromatin structure remained well-preserved, as evidenced by the integrity of nucleosomes visible in the denoised tomograms acquired from such samples (D).

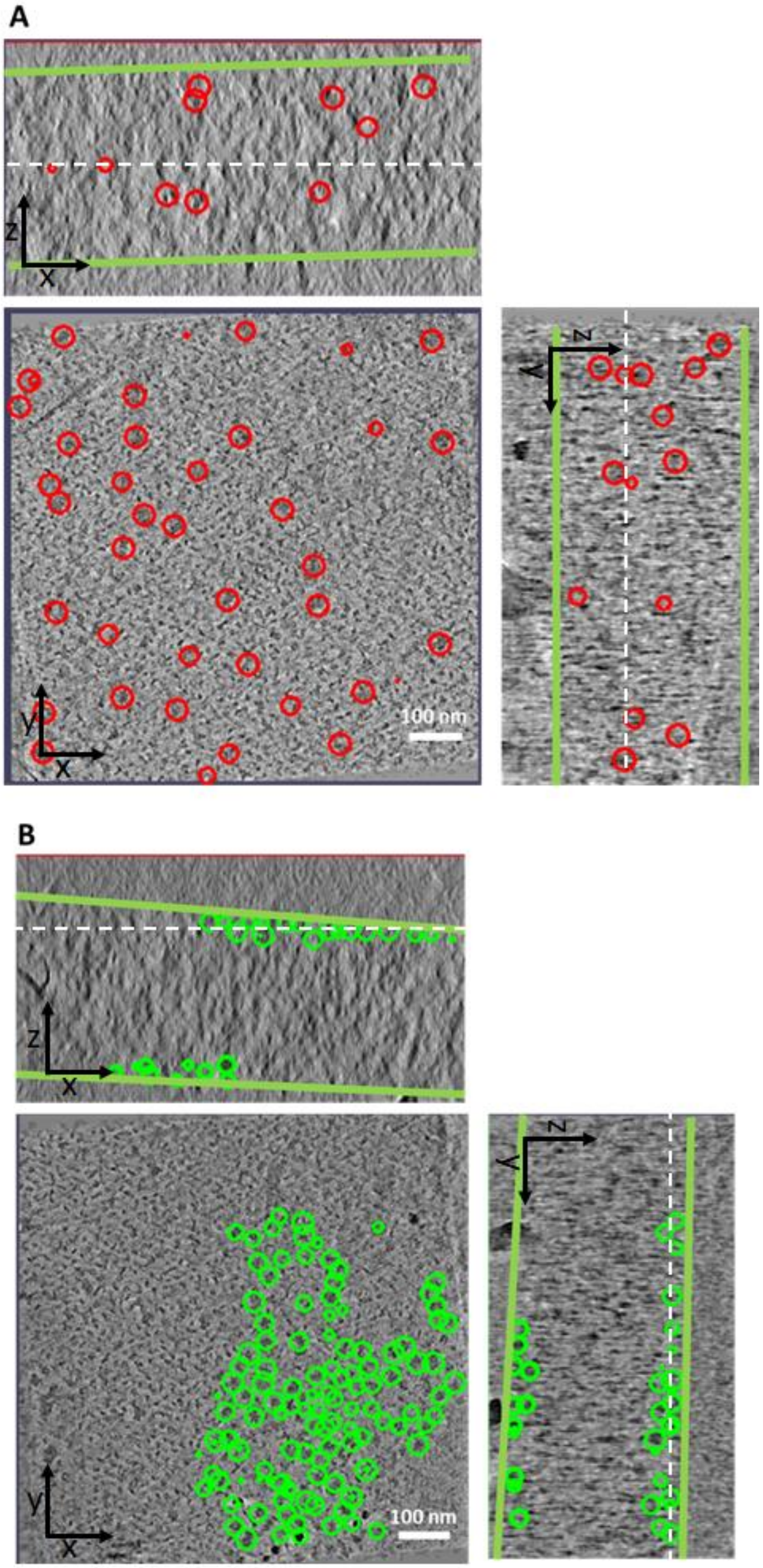

A
z
x
y
z
y
x
100 nm
B
z
x
z
y
y
x
100 nm

**Supplementary Figure 3.** (A) XYZ window of Figure 4B with manually annotated HNMDs (in red) that were observed inside the volume of the axial region of the chromosome and segmented surface of the chromosome (in light green). (B) XYZ window of Figure 5A with manually annotated surface densities (in green) and segmented surface of the chromosome (in light green). The dashed lines in XZ and YZ views show the position of the XY tomographic slice. Soft green lines mark chromosome surfaces.

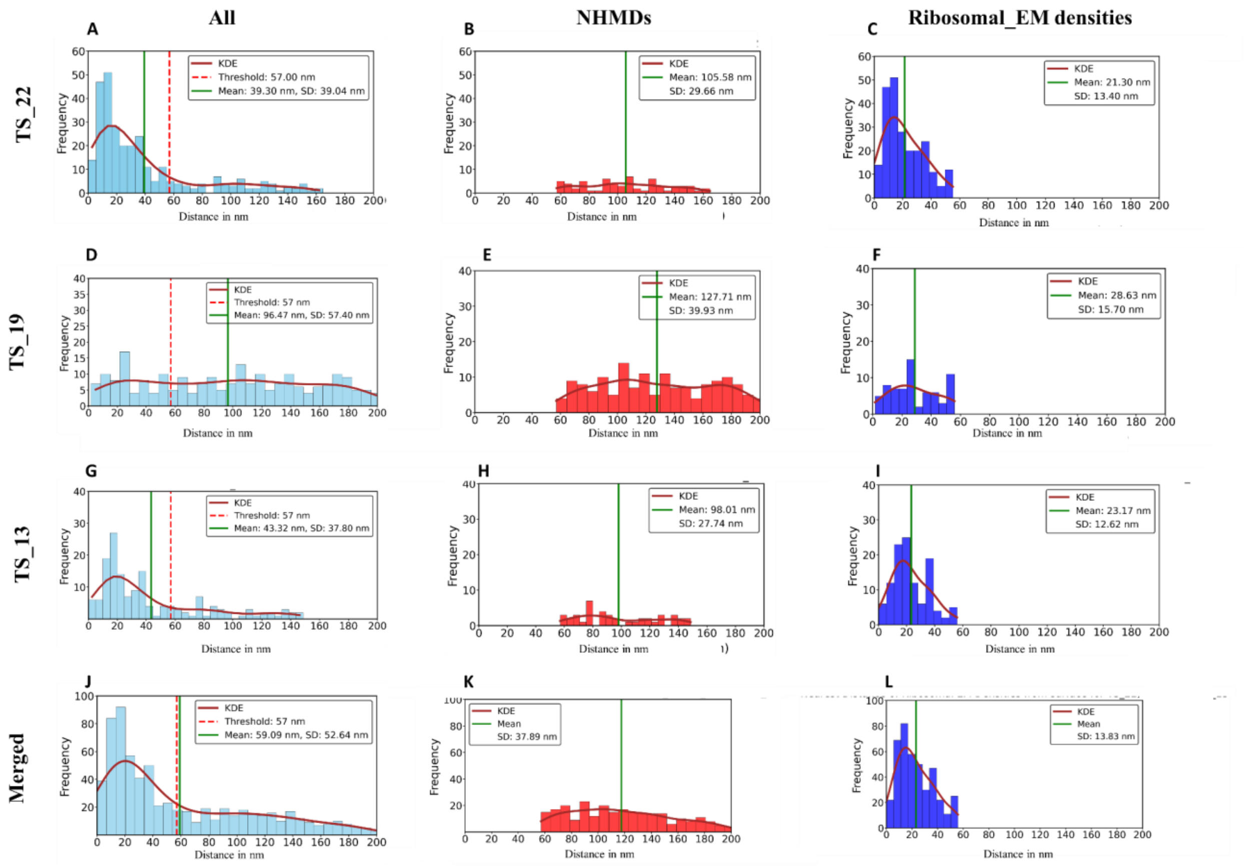


**Supplementary Figure 4:** Nearest surface distance distributions of modelled densities in TS_22, TS_19, and TS_13, showing the complete population of particles (all), surface fractions (the ribosome-related EM densities) and inner fractions (the NHMDs). The fitted smoothened curves represent the Gaussian plot. The dashed red lines represent the threshold values determined by GMM. The solid green line corresponds to the average value.

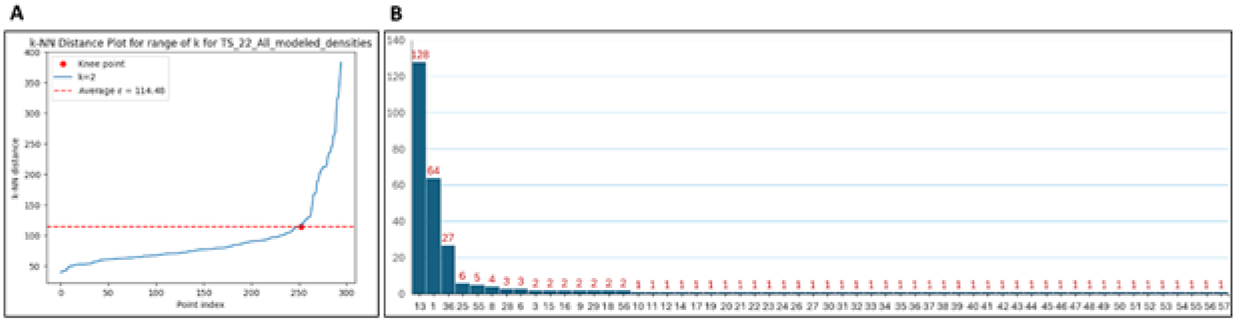


**Supplementary Figure 5:** An example of clustering analysis of particles using DBScan in the case of TS_22. (A) k-NN distance plot of all the modelled points in tomogram TS_22. (B) DBScan clustering analysis of all the modelled non-nucleosomal densities showed 57 clusters in 3D, with cluster numbers 1, 13 and 36 having the largest number of points. The model containing points belonging to these classes and its overlap with the results of the closest surface distance analysis are shown in Figure 6A.

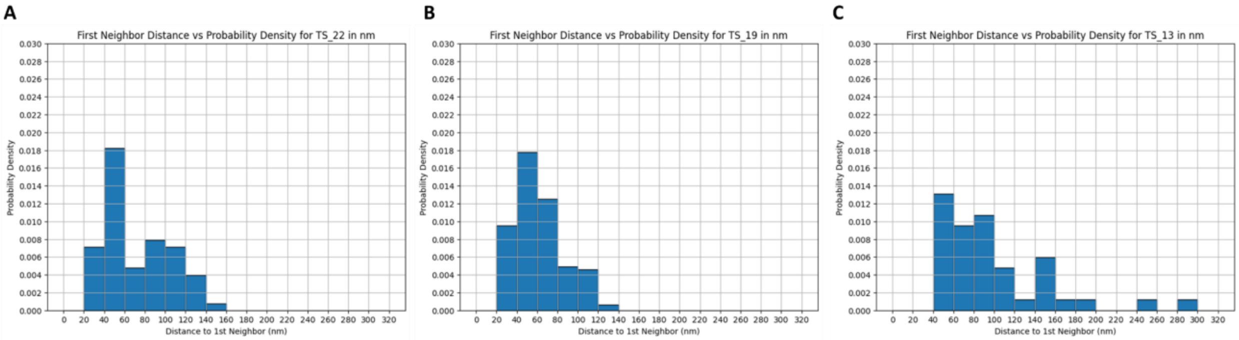


D

| Tomogram | Average distance to the first neighbour (in nm) |
|---|---|
| TS_22 | 69.94 |
| TS_19 | 62.18 |
| TS_13 | 97.64 |

**Supplementary Figure 6:** First neighbour distance analysis with an estimated probability of the NHMDs in the axial region of mitotic chromosome preparation. (A-C) First neighbour distance analysis comparing the probability of separation of the first neighbour for TS_22, TS_19 and TS_13 respectively. (D) Table showing the average distance to the first neighbour for the NHMDs in the axial region of the chromosome.

### Distribution of the nearest surface distances of the NHMDs and Ribosomal_EM densities

**A**

| | Mean (in nm) | Standard Deviation (in nm) | Skewness | Kurtosis |
|---|---|---|---|---|
| TS_22, All modelled densities | 39.30 | 39.04 | 1.50 | 1.24 |
| TS_22 NHMDs | 105.58 | 29.66 | 0.09 | -0.99 |
| TS_22 Ribosomal_EM densities | 21.30 | 13.40 | 0.69 | -0.41 |

**B**

| | Mean (in nm) | Standard Deviation (in nm) | Skewness | Kurtosis |
|---|---|---|---|---|
| TS_19, All modelled densities | 96.47 | 57.40 | 0.06 | -1.20 |
| TS_19 NHMDs | 127.71 | 39.93 | 0.07 | -1.14 |
| TS_19 Ribosomal_EM densities | 28.63 | 15.7 | 0.24 | -1.09 |

**C**

| | Mean (in nm) | Standard Deviation (in nm) | Skewness | Kurtosis |
|---|---|---|---|---|
| TS_13, All modelled densities | 43.32 | 37.80 | 1.25 | 0.51 |
| TS_13 NHMDs | 98.01 | 27.24 | 0.31 | -1.21 |
| TS_13 Ribosomal_EM densities | 23.17 | 12.62 | 0.62 | -0.228 |

**D**

| | Mean (in nm) | Standard Deviation (in nm) | Skewness | Kurtosis |
|---|---|---|---|---|
| TS_22, TS_19 and TS_13, All modelled densities | 59.09 | 52.64 | 0.94 | -0.30 |
| TS_22, TS_19 and TS_13 NHMDs | 117.43 | 37.90 | 0.62 | -0.50 |
| TS_22, TS_19 and TS_13 Ribosomal_EM densities | 23.04 | 13.83 | 0.62 | -0.502 |

**Supplementary Table 1:** (A-D) Parameters of the nearest surface distance distributions of modelled densities in TS_22, TS_19, TS_13 and all the tomograms together, showing the complete population of particles (all), and separated surface (Ribosomal_EM densities) and inner fractions (NHMDs).

**K-S two-sample test of nearest distances from the surface for NHMDs and Ribosomal_EM densities from the surface for TS_22 and TS_19**

**A**

| | TS_22 NHMDs | TS_22 Ribosomal_EM densities | TS_19 NHMDs | TS_19 Ribosomal_EM densities |
|---|---|---|---|---|
| TS_22 NHMDs | 1 | $1.27X10^{-65}$ | 0.0014 | $3.19X10^{-39}$ |
| TS_22 Ribosomal_EM densities | $1.27X10^{-65}$ | 1 | $5.4X10^{-111}$ | 0.0014 |
| TS_19 NHMDs | 0.0014 | $5.4X10^{-111}$ | 1 | $2.80X10^{-59}$ |
| TS_19 Ribosomal_EM densities | $3.19X10^{-39}$ | 0.0014 | $2.80X10^{-59}$ | 1 |

**K-S two-sample test of nearest distances from the surface for NHMDs and Ribosomal_EM densities from the surface for TS_19 and TS_13**

**B**

| | TS_19 NHMDs | TS_19 Ribosomal_EM densities | TS_13 NHMDs | TS_13 Ribosomal_EM densities |
|---|---|---|---|---|
| TS_19 NHMDs | 1 | $2.80X10^{-59}$ | 0.00072 | $5.20X10^{-78}$ |
| TS_19 Ribosomal_EM densities | $2.80X10^{-59}$ | 1 | $1.70X10^{-31}$ | 0.013 |
| TS_13 NHMDs | 0.00072 | $1.70X10^{-31}$ | 1 | $9.57X10^{-39}$ |
| TS_13 Ribosomal_EM densities | $5.20X10^{-78}$ | 0.013 | $9.57X10^{-39}$ | 1 |

**K-S two-sample test of nearest distances from the surface for NHMDs and Ribosomal_EM densities from the surface for TS_22 and TS_13**

**C**

| | TS_22 NHMDs | TS_22 Ribosomal_EM densities | TS_13 NHMDs | TS_13 Ribosomal_EM densities |
|---|---|---|---|---|
| TS_22 NHMDs | 1 | $1.27X10^{-65}$ | 0.179 | $2.88X10^{-49}$ |
| TS_22 Ribosomal_EM densities | $1.27X10^{-65}$ | 1 | $3.18X10^{-50}$ | 0.014 |
| TS_13 NHMDs | 0.179 | $3.18X10^{-50}$ | 1 | $9.57X10^{-39}$ |
| TS_13 Ribosomal_EM densities | $2.88X10^{-49}$ | 0.014 | $9.57X10^{-39}$ | 1 |

**Supplementary Table 2:** (A-C) P-values of the "Kolmogorov–Smirnov" two-sample tests comparing distributions between tomograms.
